# Individual Variation in Brain Network Topography is Linked to Schizophrenia Symptomatology

**DOI:** 10.1101/692186

**Authors:** Uzma Nawaz, Ivy Lee, Adam Beermann, Shaun Eack, Matcheri Keshavan, Roscoe Brady

**Author notes:** These authors contributed equally to this manuscript (Uzma Nawaz, Ivy Lee). Corresponding author Address correspondence to: Roscoe Brady Jr., 75 Fenwood Road, Boston, MA, 02115. This work is not peer reviewed.

## Abstract

**Background:** Resting state fMRI (rsfMRI) demonstrates that the brain is organized into distributed networks. Numerous studies have examined links between psychiatric symptomatology and network functional connectivity. Traditional rsfMRI analyses assume that the spatial organization of networks is invariant between individuals. This dogma has recently been overturned by the demonstration that networks show significant variation between individuals. We tested the hypothesis that previously observed relationships between schizophrenia negative symptom severity and network connectivity are actually due to individual differences in network spatial organization.

**Methods:** 44 participants diagnosed with schizophrenia underwent rsfMRI scans and clinical assessments. A multivariate pattern analysis determined how whole brain functional connectivity correlates with negative symptom severity at the individual voxel level.

**Results:** Brain connectivity to a region of the right dorso-lateral pre-frontal cortex correlates with negative symptom severity. This finding results from individual differences in the topographic distribution of two networks: the default mode network (DMN) and the task positive network (TPN). Both networks demonstrate strong (r∼0.49) and significant (p<0.001) relationships between topography and symptom severity. For individuals with low symptom severity, this critical region is part of the DMN. In highly symptomatic individuals, this region is part of the TPN.

**Conclusion:** Previously overlooked individual variation in brain organization is tightly linked to differences in schizophrenia symptom severity. Recognizing critical links between network topography and pathological symptomology may identify key circuits that underlie cognitive and behavioral phenotypes. Individual variation in network topography likely guides different responses to clinical interventions that rely on anatomical targeting (e.g. TMS).

## Introduction

The brain is organized into distributed resting-state networks whose spatial distribution are typically visualized using group averaging_1, 2_. Individual variation in network topography among healthy control participants has been demonstrated previously_3-8_. Until very recently, the relevance of individual network topography to cognitive and behavioral variation has been undetermined but recent reports have linked individual topography to several cognitive and behavioral phenotypes_9-11_. To date these reports have examined these phenotypes in healthy populations_9, 10_ or have examined normative processes (e.g. social cognition) in clinical populations_11_. However, the following question has been left unaddressed: Is individual network topography linked to clearly pathological (i.e. disease-specific) processes such as psychiatric symptomatology?

A very provocative idea espoused by Bijsterbosch et al. is that previously reported cross-subject variation in functional connectivity are mis-interpretations of variation in the spatial organization of brain networks_9_. We therefore sought to simultaneously address two aims: First, to determine if psychiatric symptomatology is linked to brain network topography, and second, to re-examine previously reported functional connectivity findings to determine if these findings are instead explained by individual topographic variation.

In a previous manuscript we used an unbiased, data-driven approach to find functional MRI correlates of negative symptom severity_12_. Although psychotic symptoms (e.g. hallucinations, delusions) are the most readily recognizable symptoms of schizophrenia, they are not the best predictors of functional status (e.g. independent housing and employment). Rather, the severity of negative symptoms such as amotivation, expressive deficits, and anhedonia best predict functional outcomes and overall quality of life of individuals with schizophrenia_13-15_. In our prior manuscript, we observed that the most significant resting state functional connectivity correlate of negative symptom severity was connectivity between the bilateral dorsolateral prefrontal cortex and the Default Mode Network (DMN). Here we re-examine this result in light of newly discovered relationships between functional connectivity and network topography. We examined whether negative symptom severity is linked to individual network topography. Specifically, we looked at the spatial organization of two distributed brain networks in relation to a critical region in the DLPFC.

## Methods

### Participants

Participants included in the following analysis were recruited for an Institutional Review Board approved clinical trial (NCT01561859) at Beth Israel Deaconess Medical Center (Boston, MA) and University of Pittsburg (Pittsburg, PA). Prior to enrollment in this longitudinal, randomized study, all participants gave their written informed consent. The data included in the analysis were collected at baseline (i.e. prior to trial intervention). Clinical and imaging data from 80 participants with schizophrenia were analyzed. After selection for very low-motion scans, a total of 44 participants were retained for further analysis.

Diagnosis was determined using the Structured Clinical Interview for the DSM-IV (SCID)_16_. Participants were assessed by raters trained in the administration and scoring of the SCID. Inclusion criteria for all participants were: (1) age 18-45 years; (2) current IQ ≥ 80 as assessed using the WASI-II_17_; and (3) the ability to read (sixth grade level or higher) and speak fluent English. Additional inclusion criteria for participants with a psychotic disorder were (1) a diagnosis of schizophrenia or schizoaffective disorder verified using the SCID interview_18_; (2) time since first psychotic symptom of ≤ 8 years; and (3) clinically stabilized on antipsychotic medication (assessed via SCID in consensus conferences). Healthy comparison (HC) participants did not meet criteria for any Axis I psychiatric disorder currently or historically.

Exclusion criteria were (1) significant neurological or medical disorders that may produce cognitive impairment (e.g., seizure disorder, traumatic brain injury); (2) persistent suicidal or homicidal behavior; (3) a recent history of substance abuse or dependence (within the past 3 months); (4) any MRI contraindications and (5) decisional incapacity requiring a guardian.

Demographic and clinical data for participants included in the analysis can be found in Table 1.

**Table 1.**
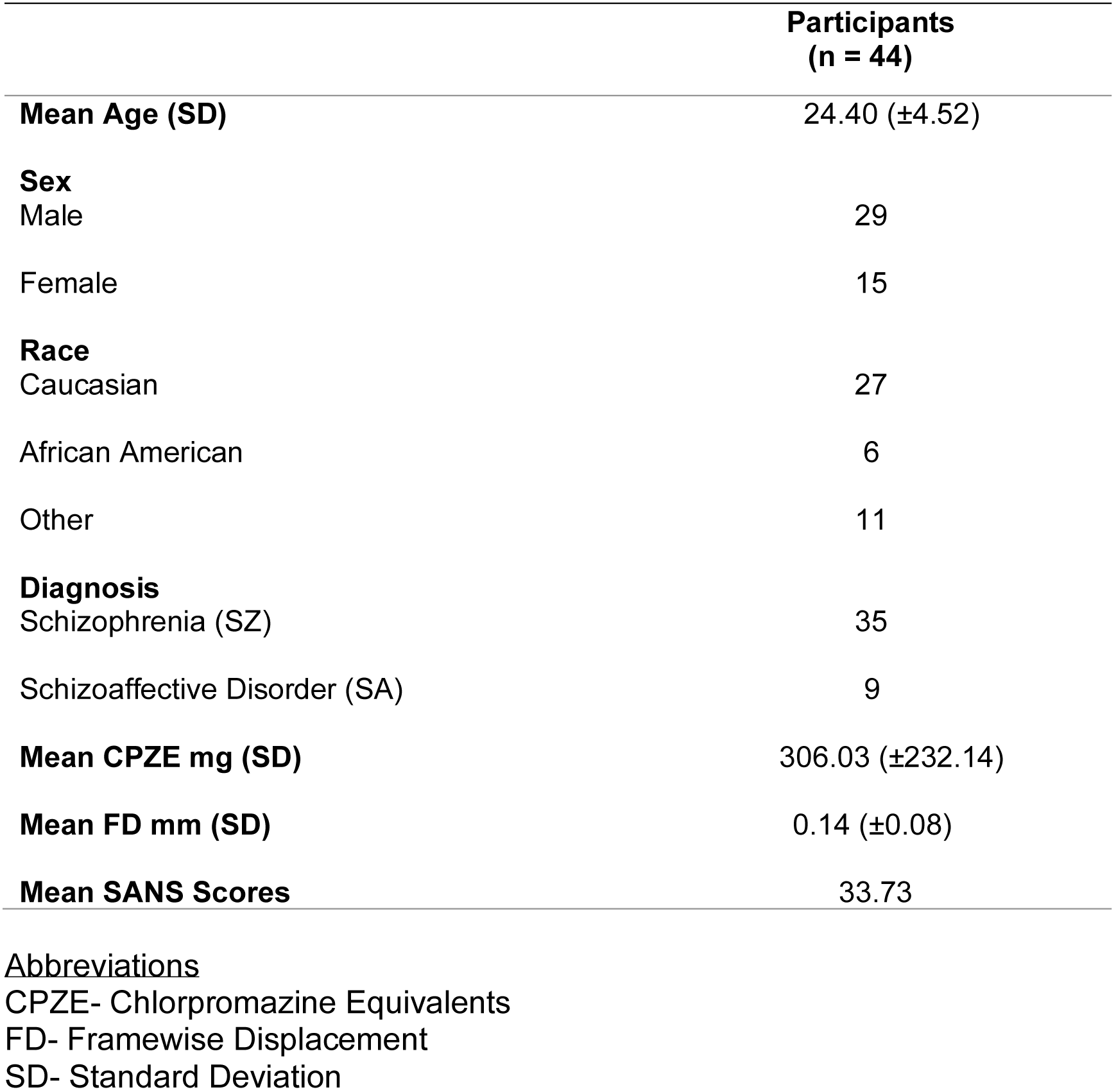
Demographic and Clinical Information for Participants.

### Behavioral Assessment

Negative symptom severity was assessed by the Scale for the Assessment of Negative Symptoms (SANS)_19_.

### MRI Data Collection

#### Boston site

Data were acquired on 3T Siemens Trio (TIM upgrade) scanners using a standard head coil. The echoplanar imaging parameters were as follows: repetition time, 3000 milliseconds; echo time, 30 milliseconds; flip angle, 85°; 3 × 3 × 3-mm voxels; and 47 axial sections collected with interleaved acquisition and no gap. Structural data included a high-resolution T1 image. In addition, all participants underwent a resting fMRI run with the instructions ‘remain still, stay awake, and keep your eyes open’. Each functional run lasted 6.2 minutes (124 time points).

#### Pittsburgh site

Data were acquired on a 3T Siemens Verio scanner using a standard head coil. The echoplanar imaging parameters were as follows: repetition time, 3000 milliseconds; echo time, 30 milliseconds; flip angle, 85°; 3 × 3 × 3-mm voxels; and 45 axial sections collected with interleaved acquisition and no gap. Structural data included a high-resolution T1 image (voxel size of 1.0 × 1.0 × 1.2 mm, TR 2300 ms, TI = 900 ms, TE = 2.89 ms, flip angle = 90°, FOV = 256 mm, 256 × 256 matrix, 160 slices, slice thickness = 1.2 mm). In addition, all participants underwent a resting fMRI run that lasted 6.2 minutes (124 time points).

### Functional MRI Data Pre-Processing

The rsfMRI data was processed using DPABI_20_. The first 4 images were removed to reduce the effects of scanner signal stabilization. Scans with head motion exceeding 3 mm or 3° of maximum rotation through the resting-state run were discarded. Functional and structural images were co-registered. Structural images were then normalized and segmented into gray, white and CSF partitions using the DARTEL technique_21_. A Friston 24-parameter model_22_ was used to regress out head motion effects from the realigned data. CSF and white matter signals, global signal and, the linear trend were also regressed out. The decision to regress the global signal was based on prior demonstration that the combination of this step plus volume-wise ‘scrubbing’ for head ‘micromovements’ is an effective strategy for removal of motion artifacts_23_.

After realigning, slice timing correction, and co-registration, framewise displacement (FD) was calculated for all resting state volumes_24_. All volumes within a scan with a FD greater than 0.2-mm were censored. Any scan with > 50% of volumes requiring censoring was discarded. After nuisance covariate regression, the resultant data were band-pass filtered to select low frequency (0.01 – 0.08Hz) signals. Filtered data were normalized by DARTEL into MNI space and then smoothed by a Gaussian kernel of 8mm_3_ full-width at half maximum (FWHM). Voxels within a gray matter mask were used for further analyses.

After preprocessing, 36 scans were removed from analysis on the basis of head motion leaving a total of 44 participants with schizophrenia or schizoaffective disorder across all sites in the study (Table 1).

### Imaging Analysis

#### Multivariate Distance Matrix Regression

We performed a connectome-wide association study using multivariate distance matrix regression (MDMR) as originally described in Shehzad et al._25_ Briefly, MDMR tests every voxel to determine if whole-brain connectivity to that voxel is more similar in individuals with a similar SANS score than in individuals with a dissimilar SANS score. A step-wise description of MDMR can be found in Supplementary Material.

This analysis identifies anatomical regions where SANS score is significantly related to functional connectivity. At each brain voxel, MDMR calculates a distance metric *r* between subjects by first calculating the temporal correlation in BOLD signal between that voxel and every other brain voxel. The generated set of temporal correlations is then correlated with the analogous set of temporal correlations from a different subject. The resulting between-subject correlation determines a distance metric that is a measure of between-subject similarity (or dissimilarity) in terms of functional connectivity at that voxel. Notably, this process disregards spatial information about the voxels that gave rise to between-individual distances. For example, two individuals may be very distant (dissimilar) in the functional connectivity of a voxel in the precuneus, but is their dissimilarity driven by differences in precuneus connectivity to the mPFC or temporal lobe or parietal lobe or all of the above? MDMR as implemented by Shehzad et al. does not display this information_25_.

To visualize this spatial information requires follow-up seed-based connectivity analysis. Shehzad et al. and others have termed this follow-up analysis ‘post-hoc’ testing to make clear that this should not be considered independent hypothesis testing nor should it be considered independent validation of the original MDMR finding_25-28_. As in these prior studies, we conducted the MDMR analysis to find anatomical regions where connectivity significantly varied with SANS score and then conducted follow-up seed-based connectivity analysis to examine the spatial distribution of these connectivity differences.

### ROI based connectivity Analysis

Following the creation and analysis of distance maps in MDMR, DPABI was used to extract the time course of the BOLD signals from rsfMRI scans in reference to a specific ROI identified in the MDMR process. Whole brain connectivity maps were generated and the resulting z-transformed Pearson’s correlation coefficients were used in the program, SPM12 (“SPM - Statistical Parametric Mapping,” http://www.fil.ion.ucl.ac.uk/spm). For the ROI generated from MDMR above, we regressed the maps against SANS score to generate spatial maps of how whole brain functional connectivity to the ROI varies with SANS scores. SPM generated T contrasts maps were created using Caret_31_. Correlations between ROI to ROI connectivity values and SANS score were calculated in R.

## Results

### Resting State Connectivity to the Bilateral DLPFC Is Correlated to Negative Symptom Severity

From the MDMR analysis of 44 participants, two regions were identified where resting state functional connectivity correlated to SANS scores (Figure 1). Functional connectivity to the bilateral middle frontal gyrus (Brodmann Area 9) in the dorsolateral prefrontal cortex covaried significantly with negative symptom severity. The right dorsolateral prefrontal cortex (peak voxel z-stat 3.82 at MNI coordinates: x36, y24, z30) demonstrated a more significant relationship between functional connectivity and negative symptom severity than the left (peak voxel z-stat 2.95 at MNI coordinates: x-33, y30, z42) (Figure 1F).

**Figure 1.**
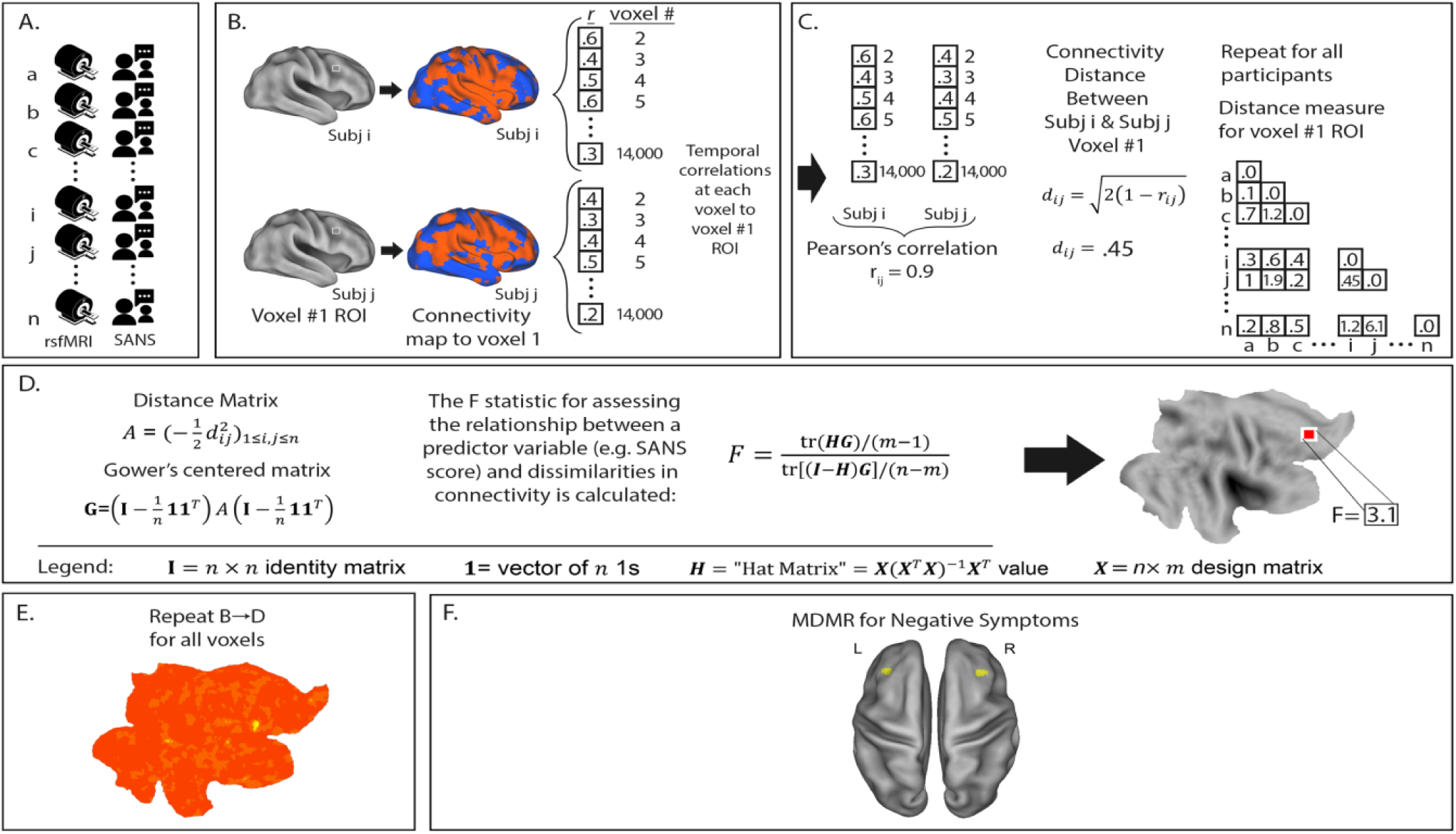
Multivariate Distance Matrix Regression identifies brain voxels whose functional connectivity varies with negative symptom severity. MDMR procedure: **A**. rsfMRI and SANS are collected from each participant. **B**. For each participant a functional connectivity map is generated to an individual voxel. **C**. Voxelwise temporal correlations between participants are used to generate a Pearson’s correlation *r* and a distance metric *d*. This is repeated for all participants to generate a matrix of between subject distances. **D**. The distance matrix is centered and an ANOVA-like test is used to generate an F-statistic to assess the relationship between a predictor variable (SANS score) and dissimilarities in functional connectivity at that voxel. **E**. This process is repeated for every voxel. This results in a whole brain map of how significantly functional connectivity is related to emotional intelligence. Permutation testing then identifies whole-brain significant clusters in connectivity-SANS score relationships. **F**. In our sample of 44 participants with schizophrenia or schizoaffective disorder, we identified the bilateral DLPFC (Brodmann Area 9) as regions where resting-state connectivity correlated significantly with SANS score. In this image, connectivity is thresholded at a voxelwise level of p<.005 and extent threshold of p<.05.

### The Spatial Distribution of Symptom-Correlated Connectivity to the DLPFC Identifies the Default Mode Network and Task-Positive Network

We performed follow-up analysis using the right DLPFC in a seed-based connectivity analysis to determine the spatial distribution and directionality of connectivity that gave rise to this result. This analysis revealed two apparent patterns of functional connectivity: higher SANS (i.e. more severe symptoms) correlated with increasing intrinsic connectivity between the right DLPFC and the “task-positive network” (TPN) (Figure 2A)_32_. Inversely, lower negative symptom severity correlated with increasing functional connectivity between the right DLPFC and the “default mode network” (DMN) (Figure 2B)_1_.

**Figure 2.**
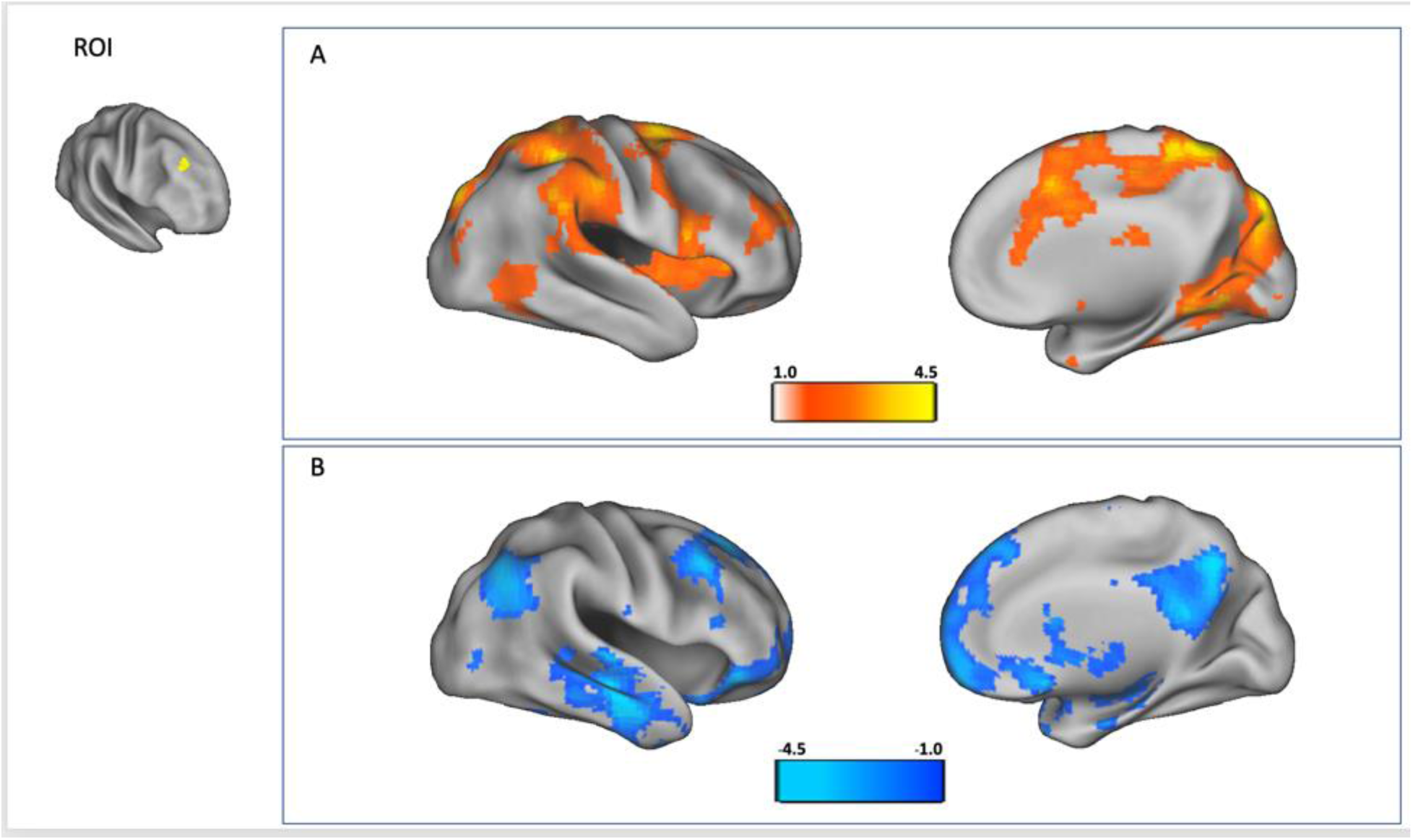
In a follow-up analysis, a seed-region was placed in the right dorsolateral prefrontal cortex (ROI) in all subjects and then this seed-based connectivity map was correlated with SANS scores to identify locations where increasing connectivity to dorsolateral prefrontal cortex corresponds to more severe symptoms (red) and decreased connectivity corresponds to less severe symptoms (blue). Here, we observe that right DLPFC connectivity to the Task-Positive Network (TPN) is correlated with higher SANS score (A) and right DLPFC connectivity to the Default Mode Network (DMN) is correlated with lower SANS score (B). Color bar = T-statistic.

### Individual Variation in Network Topography Explains Connectivity-Symptom Relationships in the DLPFC

We plotted the relationship between negative symptom severity and functional connectivity for both networks as a scatter plot in Figure 3C & 3D. As expected from the maps in Figure 2A & 2B, we observed that a higher SANS score was correlated with increased functional connectivity between the right DLPFC and the TPN and decreased connectivity to the DMN. Both scatter plots of connectivity cross the axis of 0 connectivity i.e. activity in this DLPFC region is intrinsically correlated to the DMN in some individuals and is intrinsically correlated to the TPN in others. Furthermore, this network membership is organized along an axis of negative symptom severity with this critical zone included in the spatial distribution of the DMN in the least symptomatic individuals and, in the most symptomatic individuals this region is included in the TPN.

**Figure 3.**
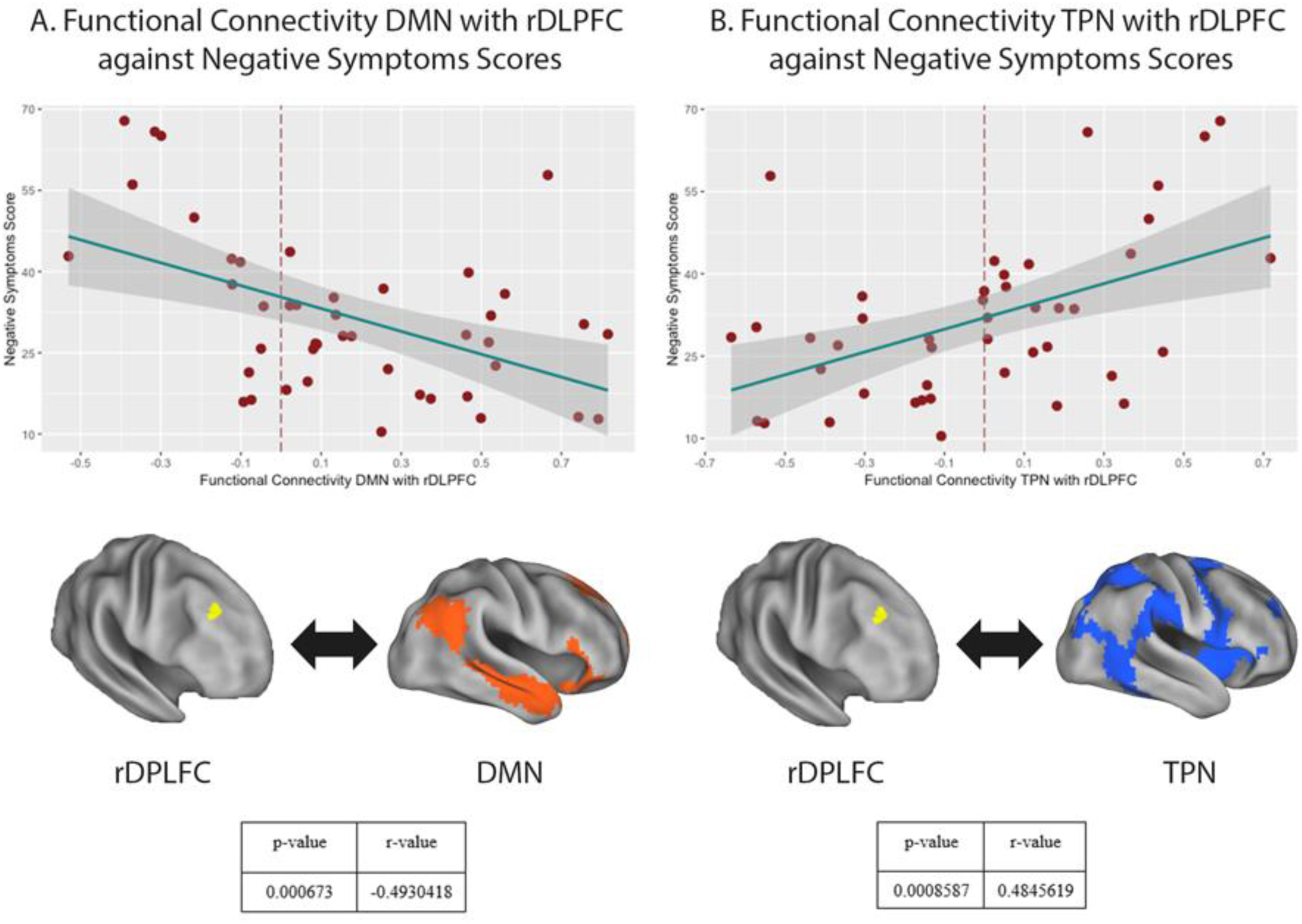
Negative Symptom Severity is Linked to Network Topography. **A**. Plot of SANS score (y-axis) and functional connectivity between the rDLPFC ROI and the DMN network. **B**. Plot of SANS score (y-axis) and functional connectivity between the rDLPFC and TPN network. Notably, both graphs demonstrate correlations that cross the reference line of null functional connectivity, meaning that the network association of this region appears to shift from being a member of the DMN to a member of the TPN as one moves along the distribution from less symptomatic individuals to more symptomatic individuals. Network topography correlates were highly significant (p<.001) for both networks.

We sought to re-visualize the relationship between symptom severity, the critical right DLPFC region, and the whole-brain spatial organization of the DMN. To facilitate this, participants were grouped by symptom severity. Forty-four participants were split into three groups based on their SANS score with group 1 consisting of the lowest scorers (i.e. least symptomatic) and 3 with the highest scores (most symptomatic). We examined the spatial distribution of the DMN in each group and visualized this in relation to the previously identified right DLPFC (Figure 4). As would be expected from Figure 3, we observe the spatial extent of the DMN does not include the identified DLPFC node in the most symptomatic participants. The DLPFC node of the DMN is organized along an anterior-posterior spatial gradient organized along a least symptomatic to most symptomatic axis. The spatial extent of the TPN (data not shown) demonstrated the inverse relationship i.e. the TPN extending into the DLPFC critical region in the most symptomatic participants.

**Figure 4.**
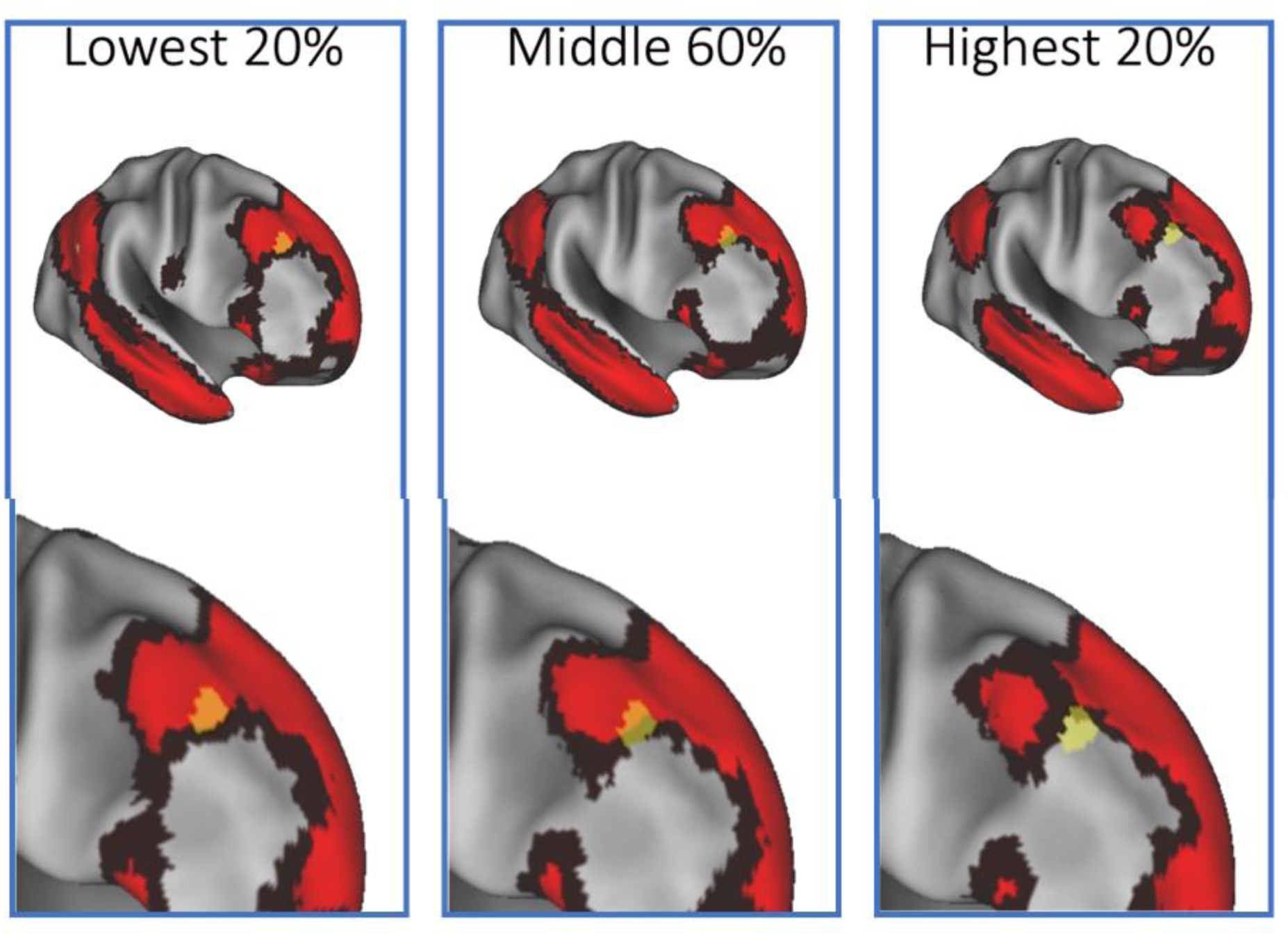
Whole Brain DMN Spatial Topography in Relation to Negative Symptom Severity. Shown is the distribution of DMN correlated voxels in the participants grouped by SANS score i.e. lowest (least symptomatic) 20%, middle 60% and highest (most symptomatic) 20% of the sample. In least symptomatic participants the rDLPFC was part of the DMN. In the most symptomatic (Group 3), this region was a part of the TPN. Groups with intermediate severity were intermediate in their connectivity strength to the DMN. Scale: Connectivity strengths (Pearson’s *r*) less than 0.25 are shown in dark red. Higher strengths in bright red. The MDMR identified right DLPFC region is shown in yellow.

## Discussion

We conducted an entirely data-driven connectome wide study examining the relationship between negative symptoms severity and functional connectivity. Bilateral dorsolateral prefrontal cortex regions were identified as having the most significant relationship between functional connectivity and psychopathology. Follow-up testing demonstrated that this result is driven by the correlation between DLPFC activity and two distributed brain networks: the Default Mode Network (DMN) and the Task Positive Network (TPN).

Analyses that assume inter-individual homogeneity in the spatial distribution of these networks would misinterpret this result as the product of individual variation in the temporal correlation in activity between these networks and the DLPFC. Instead, we observe that this result is entirely driven by individual variation in spatial organization (i.e. network topography) within this region.

More specifically, the degree to which this region was a member of one of two large scale resting-state networks, the DMN and TPN, was directly linked to symptom severity.

The implication of this result is that recently observed links between variation in network topography and normative cognitive / behavioral phenotypes also apply to disease-specific symptoms_9-11_. More broadly, this result lends support to the hypothesis proposed by by Bijsterbosch et al. that previously reported cross-subject variations in functional connectivity are in fact misinterpretations of variation in the spatial organization of brain networks_9_.

It is notable that in both this manuscript and our previous report on social cognition_11_, we demonstrated that the *most significant* connectivity-phenotype relationships in the entire brain were driven by variation in network topography. The increasing prevalence of fully data-driven connectivity studies will determine if this (i.e. spatial variation driving the most significant connectivity-phenotype relationships) is a general rule.

Returning to this specific result, what are the implications of schizophrenia symptoms being linked to network organization within a narrowly circumscribed portion of the DLPFC? The DLPFC has been implicated in an enormous variety of cognitive processes and thus links between DLPFC organization and negative symptomatology have face validity. From a pragmatic/experimental perspective, our finding has clearer significance: An enormous body of literature has explored the activation of the DLPFC in a variety of neuroimaging tasks. Although our result was generated using task-free connectivity data, the at-rest spatial organization of the TPN would be expected to represent the spatial organization of the networks activated under task conditions. The implication of our finding is that there is considerable variability in the organization of this network in a population diagnosed with schizophrenia. This would almost certainly confound task-based studies that rely on group-based comparisons. Put simply: In participants with schizophrenia, the DLPFC region activated by a task may lie in very different MNI coordinates even after normalization to a standard template. This result would likely be interpreted as hypoactivation simply because of a lack of overlap among a group when compared to a groups of healthy comparison participants.

Our results also have clear implications for therapeutic interventions that rely on the precise spatial targeting. In a previous publication, we demonstrated that repeated application of intermittent Theta Burst Stimulation (iTBS) targeted to the cerebellum can restore connectivity between cerebellar and DLPFC nodes of the DMN and, in the process, can reduce negative symptom severity_12_. In contrast, one of the largest studies of rTMS applied to the left DLPFC for negative symptom amelioration did not observe a significant difference between sham and real rTMS_33_. A re-analysis of that trial demonstrated that response in the real TMS group was linked to within-trial change in brain structure with responders showing an increase in the volume of DMN-related regions including hippocampus, parahippocampus, and precuneus_34_.

We hypothesize that this result is directly linked to our network topography result in the following manner: In Wobrock et al., rTMS was targeted to EEG position F3. This scalp target lies directly adjacent to the left DLPFC region of substantial individual network topography variation_33_. In the Wobrock et al. trial’s selection of a participant population with high negative symptom burden, the implication of our results presented here would be that this target would overlie the DMN only in a subset of participants. We hypothesize that rTMS response was partially determined by individual DMN network topography: When the TMS coil position coincided with the topography of an individual participant’s DMN, symptom improvement and DMN network related structural changes followed.

This logic, applied broadly, may explain individual variation in response to anatomically targeted interventions in other disorders e.g. rTMS for depression. Prior claims linking group-average connectivity maps of rTMS targets to antidepressant response may in fact, represent movement of coil position from regions of high individual variation in network topography to regions where a majority of patients show shared topography of disease relevant networks_35_. This remains a conjecture as we have not analyzed differences in network topography among depressed candidates for TMS.

Finally, our work contains a cautionary note for imaging studies examining connections between connectivity and individual phenotypes. The most frequently used approaches to identifying individual-level network topography do so by generating individual parcellations of the cortex using a “winner-take-all” approach i.e. spontaneous activity in a given cortical region is frequently correlated with *multiple* networks. In a “winner-take-all” or “hard” parcellation approach, the network “identity” assigned to that region is that of the single network most strongly connected to that region. Subsequent analyses then typically analyze results in terms of connectivity to that “winner” network.

While this approach allows for easy visualization of individually parcellated connectomes, this approach *continues* to obfuscate links between network topography and phenotypes. Specifically: The region we identify in this manuscript lies at the intersection of several different networks. At the resolution used for rsfMRI imaging these networks all show varying levels correlation within the voxels identified in our DLPFC region. While the topography of two of these networks (DMN and TPN) are very significantly linked to symptomatology, a winner-take-all analysis would actually have assigned this parcel of cortex to the FPCN (Supplementary Material Figure 1). The consequence of this is likely that subsequent analysis of connectivity-phenotype relationships would be misinterpreted as inter-network connectivity i.e. FPCN-DMN and FPCN-TPN connectivity correlations with symptomatology. This is, in fact, *exactly* what was described in a recent publication (Figure 2, Baker et al._36_) which uses an individual “winner-take-all” parcellation approach in an independent dataset to elucidate connectome-negative symptom relationships.

In summary, we demonstrate that inter-individual variation in disease phenotypes is linked to individual network spatial organization. We present a model for how this variation in cortical organization is likely to explain previously observed variation in response to targeted neuromodulation. We also highlight how commonly used individual parcellation approaches continue to obfuscate significant network topography-phenotype relationships.

## Acknowledgements

Funding:

This work was supported by the National Institutes of Health (K23 MH100623 and R01 MH116170 to RB, R01 MH92440 to SE, MK)

## Supplemental Information

### Multivariate Distance Matrix Regression

As previously described ^26-28^, this analysis occurs in several stages: First, a seed-to-voxel connectivity map is generated by using an individual voxel’s BOLD signal time-course to calculate the temporal Pearson’s correlation coefficients between that voxel and all other gray matter voxels (Figure 1B). These maps are generated for every participant. In the second stage, the temporal correlation coefficients for each voxel in the connectivity map is correlated with the values of corresponding voxels in maps generated from every other participant. This Pearson’s correlation coefficient, *r* is a measure of the similarity of whole-brain connectivity to that voxel between patients. This value is used to calculate between-subject distance (or dissimilarity) using the metric 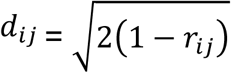 where *i* and *j* are two subjects and *r* is the correlation coefficient above. (Figure 1C)^29^. The next stage tests the relationship between a given variable (here SANS score) and the inter-subject distances in connectivity generated in the previous stage.

Broadly speaking, this process consists of ANOVA-like hypothesis testing where the tested relationship is between a variable of interest and a matrix of distances. This process was originally termed multivariate distance matrix regression by Zapala and Schork and used to test associations between gene expression and related variables ^29^. Shezhad et al. then used their framework for testing the relationship between variables of interest and a matrix of distances where the matrix is between-subject similarity in whole-brain functional connectivity.

This test consists of first forming a distance matrix 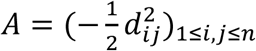 among *n* participants where *d* = the between subject distance metric calculated above. Next, this matrix is used to create a Gower’s centered matrix 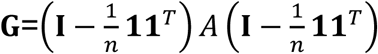, in which *n* is the number of participants, **I** is the *n* × *n* identity matrix, and **1** is a vector of *n* 1s. The *F* statistic for assessing the relationship between a predictor variable (e.g. SANS score) and dissimilarities in connectivity is calculated as follows: For *m* predictor variables, let ***X*** be a *n*× *m* design matrix of predictor values, and let ***H*** = ***X***(***X***^*T*^***X***)^−1^***X***^*T*^ be the associated *n* × *m* “hat” matrix

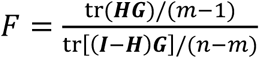 (Figure 1D) ^25^. This process is repeated for every voxel.

The result is a whole brain map of how significantly SANS scores are related to functional connectivity at every voxel (Figure 1E). Covariates included in this analysis included scanner site, age, sex, mean FD (as a subject-level correction for motion effects^24^), and prescribed antipsychotic dosage (chlorpromazine equivalents, CPZE). To correct for multiple comparisons, a nonparametric permutation was calculated for voxels that exceeded the significance threshold of p < 0.005 and clusters of such of sizes at a threshold of p < 0.05 ^30^, with a null distribution calculated from 5000 such permutations, as in prior MDMR studies.

## Supplemental Figure 1

**Supplemental Figure 1.**
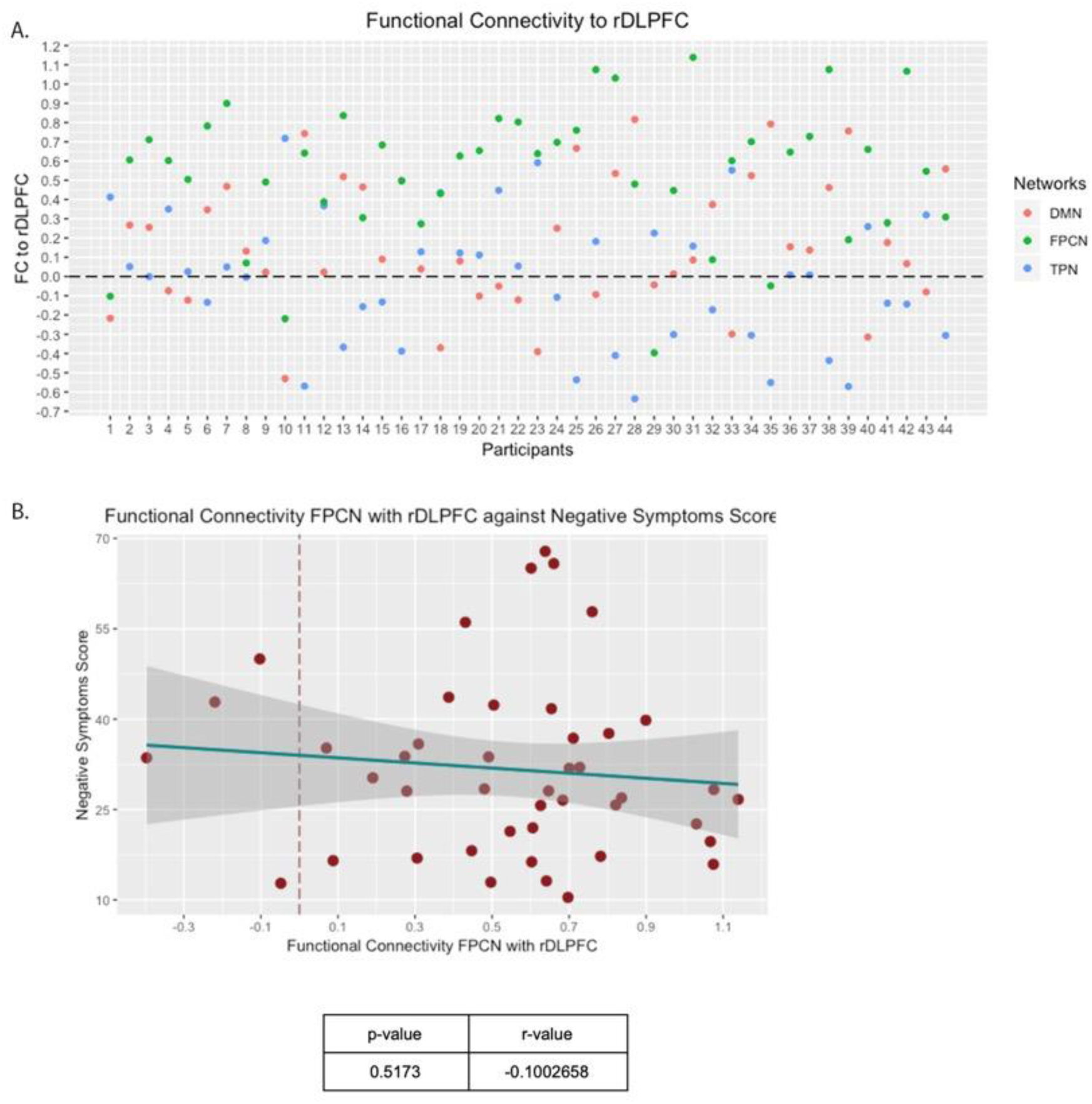
Winner-Take-All Parcellations Obfuscate Topology-Phenotype Relationships. Several approaches to individual parcellation use a “winner-take-all” approach in assigning cortical territory to one canonical resting state network or another. We explored the consequence of this approach when applied to our dataset. In A) the connectivity strength between the right DLPFC region of interest and three resting state networks (DMN, TPN and FPCN, Fronto-Parietal Control Network) were plotted for every individual in the dataset. Using a winner-take-all parcellation, this region would have been assigned to the FPCN in 75% of participants. In B) we demonstrate that within-network connectivity between the right DLPFC region and the FPCN is not correlated with symptom severity. We conclude that an analysis of connectivity-symptom relationships using a winner-take-all approach would then likely be misinterpreted as correlations between symptoms and *inter*-network connectivity i.e. between the FPCN and both the DMN and TPN when, in fact, symptom-connectivity relationships at this region can be entirely described by DMN and TPN topology irrespective of the FPCN.

